# The Landscape of Maize-Associated Bacteria and Fungi Across the United States

**DOI:** 10.1101/2023.07.11.548569

**Authors:** Corey R Schultz, Hanish Desai, Jason G Wallace

## Abstract

The maize microbiome consists of microbes that are associated with plants, and can be shaped by the host plant, the environment, and microbial partners, some of which can impact plant performance. We used a public dataset to analyze bacteria and fungi in the soil, rhizosphere, roots, and leaves of commercial maize at 30 locations across the US. We found that both tissue type and location had significant effects on community structure and makeup, although the patterns differed in bacteria and fungi based on tissue type. We also found many differences in predicted microbial gene pathways between tissues, with location also shaping predicted functional gene profiles. We found a pattern of potential interaction between fungi and bacteria, and potential intra-kingdom mutualism, in microbiome networks. The robustness of these networks was dependent upon tissue, with endophytes in leaves and roots showing significantly higher natural connectivity. Within a tissue, this connectivity was relatively stable across locations. We identified environment and soil characteristics that may impact tissue specific microbial abundance. Sulfate level in the soil was positively correlated with Proteobacteria abundance, but negatively correlated with Firmicutes abundance in the roots and leafs. Ascomycota appears to be affected by different environmental variables in each tissue. We also identified gene functions and enzymes which may be necessary to allow microbes to transition across compartments and become endophytes.

## Introduction

Communities of fungi and bacteria live in, on, and around all plants [1, 2]. These microbes can colonize surfaces of tissue above ground (the phylosphere), soil near the root surface (the rhizosphere), and inside plant tissue (the endosphere) [1, 3], and they can have a large impact on plant health [1]. A plant’s microbiome—the community of microbes affiliated with it—can benefit the plant by protecting against abiotic stress [4–8], defending against pathogens and herbivores [9–12], producing phytohormones [13–15], and promoting growth through nutrient acquisition (N fixation, P solubilization, siderophore production, etc.) [2, 16–20]. Microorganisms can jumpstart plant immune systems resulting in higher levels of protections from pathogens [21–23]. On the other side, host plants can affect microbes by alterer soil chemistry and producing signaling compounds [24–27]. Plants can also secrete energy-rich carbon compounds (“root exudates”) into the environment [25] and otherwise provide habitat for a wide range of organisms [28].

Plant genetics impact interactions with both individual beneficial microorganisms [29–32], as well as the overall microbiome community [3, 12, 33–36]. The environment provides most of the microbes for community recruitment (outside of seed endophytes [35]), and it has a direct impact on soil [37], plants [38], and maize [39] microbial community structure. Changes in the environment and abiotic stress such as drought [8, 40–42], heat [40, 43]], soil salinity [44–46], and fertilizer [47–49] have all been shown to cause changes in a plant’s microbiome. Network analysis in plant microbial communities has been widely adopted to uncover underlying correlations amongst bacteria and fungi, [50–54] and many of these correlations have been found to support antagonistic between the two kingdoms [55–57]. This analysis can identify cross-kingdom interactions, as well as identify hub taxa, taxa that are highly connected and have the potential to influence microbiome network stability (which describes the proportion of correlations amongst taxa) [58–60]. There have been few studies examining the impact of biotic and abiotic environmental factors have on cross kingdom network interactions [61–63]. Maize’s high economic value ($9.2 billion in US exports alone in 2020) [64, 65] and extensive genetic variation have made it a model organism for the last century [66]. Although many studies have looked at the maize microbiome [3, 33–36, 67, 68], the majority of these studies use inbred research lines, while few looked at normal production conditions or used commercial varieties [69–72]. As microbial interactions can be effected by host genetics, current understanding and assumption about the maize microbiome using dedicated inbred lines may not translate to high performing commercial varieties.

The Genome to Fields Initiative is a large public collaboration to expand our knowledge about Genome by Environment interactions. Innovations in phenotyping have been developed to explore how weather, soil, and management practices effect crop performance. This collaboration between crop scientists, computation scientists, and engineers spans universities, government agencies, and industry. A number of publications have emerged from this collaboration, especially in the fields of GxE and Genomic Selection [73–75], Phenotyping [76–78], and robotics [79]. Microbiome data has been collected for inbred and their hybrid maize over several years [Li et al. *in preparation*], but these cultivars are research lines which may not reflect high performance commercial maize used by producers. In 2017 Indigo Ag sampled the microbiome of commercial maize border corn in 30 locations across the US, allowing us to fill a crucial gap in current research: ’differences in the microbiome of high-performing maize across environments. This data set allows us to investigate bacterial and fungal community structure, interaction, and function in the soil, rhizosphere, roots, and leaves. In this study we used an unanalyzed public dataset of maize-associated fungi and bacteria at 30 locations across the United States to (1) characterize bacterial and fungal communities, (2) identify hub taxa and inter-kingdom interactions (3) identify the impact environment has on microbiome networks, and (4) identify the core functions of the commercial maize microbiome.

## Methods

### 2.1 Data Acquisition

To briefly describe how the dataset was generated: Plants in the V3-V5 stage as well as a trowel of soil were dug up, bagged, placed into a cooler and mailed overnight for processing. DNA was extracted from samples, and bacterial and fungal amplicons were sequenced. For bacteria, the 16S sequencing used modified 515F-806R primer pairs [80–82]. For fungi, the ITS region was amplified with the P-ITS1 and P-ITS4 primers [83]

The Genomes to Fields and Indigo Ag Microbiome Data 2017 is publicly available and was downloaded from Cyverse data commons (DOI: 10.25739/htck-sw56). Weather, soil, and field meta data was also downloaded from Cyverse data commons (DOI:10.25739/frmv-wj25). Due to lack of meta data for a large percentage of samples, supplemental weather data was queried from Visual Crossing Weather API [84]. A number of daily weather variables, including solar, temperature, and precipitation metrics, were queried from March 1^st^ 2017 – August 1^st^ 2017 and then the average was used for each location.

### 2.2 Bioinformatics

#### Amplicon Sequence Variant Calling

Amplicon sequence quality filtering and processing was performed with version 2022.11 of the QIIME2 toolbox [85]. Primers were trimmed from raw sequences, and reads below a Phred score of 20 were dropped using Cutadapt [86]. FastQC [87] was used to visually confirm read quality. Paired reads were then joined via vsearch in QIIME2 [88]. Dada2 [89] was used to trim fungal reads to 250 bp, and bacteria reads to 200 bp, and cluster ASVs. Unite version ver9_99_16.10.2022 [90] and SILVA-138-99 [91] dataframes were used with QIIME2’s classifier to assign taxonomy to fungal and bacterial ASVs, respectively. The initial (raw) dataset contains 213,990 bacteria ASVs in 604 samples, and 95,058 fungal ASVs in 583 samples. We removed ASV’s from a sample that (1) were unidentified at the phylum level, (2) were identified as mitochondria or chloroplasts, or (3) had two or fewer total reads in that sample. Samples with fewer than 500 remaining reads were dropped from the analysis. This resulted in a final data set of 6,578 ASVs from both kingdoms across 1,073 samples (with bacteria and fungi separated). For analyses looking at environmental covariates, we used a subset of 734 samples with the necessary metadata. When we combined bacteria and fungal samples from the same extraction (single tissue in a single plant), we had 496 samples that had both bacteria and fungal reads associated with it.

#### Diversity Metrics

Alpha and beta diversity were compared separately for bacteria and fungi, ASV count data was transformed into proportional data or relative abundance (total sum scaling), where each sample had a read depth of 10,000 reads. We compared alpha diversity based on tissue, location, and plant stage at harvest, using the metrics of observed ASVs, Shannon, and Simpson indices, from the phyloseq package [92]. A marginal PERMANOVA from the vegan package [93] was used for each alpha diversity metric generated in phyloseq to test the relationship between metadata variables and community diversity. The marginal PERMANOVA was used to ignore term order in the model, and analyze the model term with all other terms. Bray-Curtis (tree-independent) distance matrices were generated in phyloseq for beta diversity of fungi. These measures were plotted, and the effect metadata variables had on community diversity was tested with PERMANOVA again in vegan. We then subset our data based on tissue type, and combined samples within a single field. A mantel test in vegan was used to compare Bray-Curtis distance with geographic distance between fields.

#### Network Analysis

Co-occurrence networks were built using the NetCoMi package [94] to explore correlations of bacteria and fungi. Our dataset was subset into four tissue types, and then agglomerated to the phylum level in phyloseq. This left us with 48 phyla (28 bacteria, 1 Archaea and 19 fungal). We then created our correlation network using the following parameters: we used SPIEC-EASI [95] a correlation algorithm specifically designed for compositional data, added pseudo counts of 1 to account for taxa with 0 reads, had a rho (correlation) threshold of 0.8, and normalized the data with total sum scaling [96], also known as proportions or relative abundance. These strict parameters were used to ensure we were finding the strongest and most significant correlations amongst phyla.

We used Natural Connectivity [97] to identify the effects of tissue, location, and environmental variables had on network connectivity. Natural Connectivity is a measure of network robustness, and quantifies the proportion of hubs connected to one another. This was done to ensure we captured a more holistic view of microbiome interactions. We subset our ASV data based on samples from the same tissue in a single location (∼5 samples). Networks were created in NetCoMi with the following parameters: we used spearman correlation, with Centered log-ratio normalization method [96] as spearman is not compositionally aware. Our rho (correlation) threshold was decreased to 0.5 to capture a larger proportion of potential interactions. We tested whether environmental covariates impacted network connectivity by performing Type II ANOVA using the car package in R [98], environmental variables were centered and scaled prior to testing.

#### Taxa Correlations with Environment

Correlation of microbial phyla to environmental variables was carried out with the phylosmith package [99]. We transformed our ASV data to relative abundance, agglomerated to the phylum level, added pseudocounts of 1, and then subset by tissue. A Spearman correlation test was run on bacteria and fungi separately. To account for multiple testing, p values were adjusted using the Benjamini & Hochberg (False Discovery Rate) method [100].

#### Predicted Functional Genetics

PICRUST2’s [101] standard pipeline was used to investigate predicted functional gene pathways for bacterial and fungal communities. For 16s reads, raw KO terms were agglomerated to KEGG gene families terms [102], while ITS reads used raw Enzyme Classification [103] numbers, as support for ITS functionality is more limited. Core functions were filtered to include pathways that were abundant in 90% of a single tissue across all locations. These pathways were transformed with a variance stabilizing transformation using DEseq2 [104] in order to visualize functional gene profiles. DEseq2 was then used to identify differentially abundant functional pathways between tissues, with an alpha value of .001.

## Results

### Diversity Analysis

We used public 16S and ITS sequences from the 2017 Genomes 2 Field microbiome dataset. After quality control and filtering, our final dataset consisted of 6,578 ASVs across 1,073 samples, consisting of four tissues in 30 locations across the US. Our tissue types could also be classified as endophytes (root and leaf) or exterior (soil and rhizosphere). Throughout the experiment, we found that leaf and root tissue had fewer associated ASVs compared to root wash and soil samples. At the phylum level, soil and rhizosphere have more similar taxonomic diversity and relative abundance profiles, while root and leaf samples are more similar to each other (Figure 1). In all four tissues bacterial microbiomes are dominated by Proteobacteria reads, while fungal microbiomes are dominated by Ascomycota reads.

**Figure 1.**
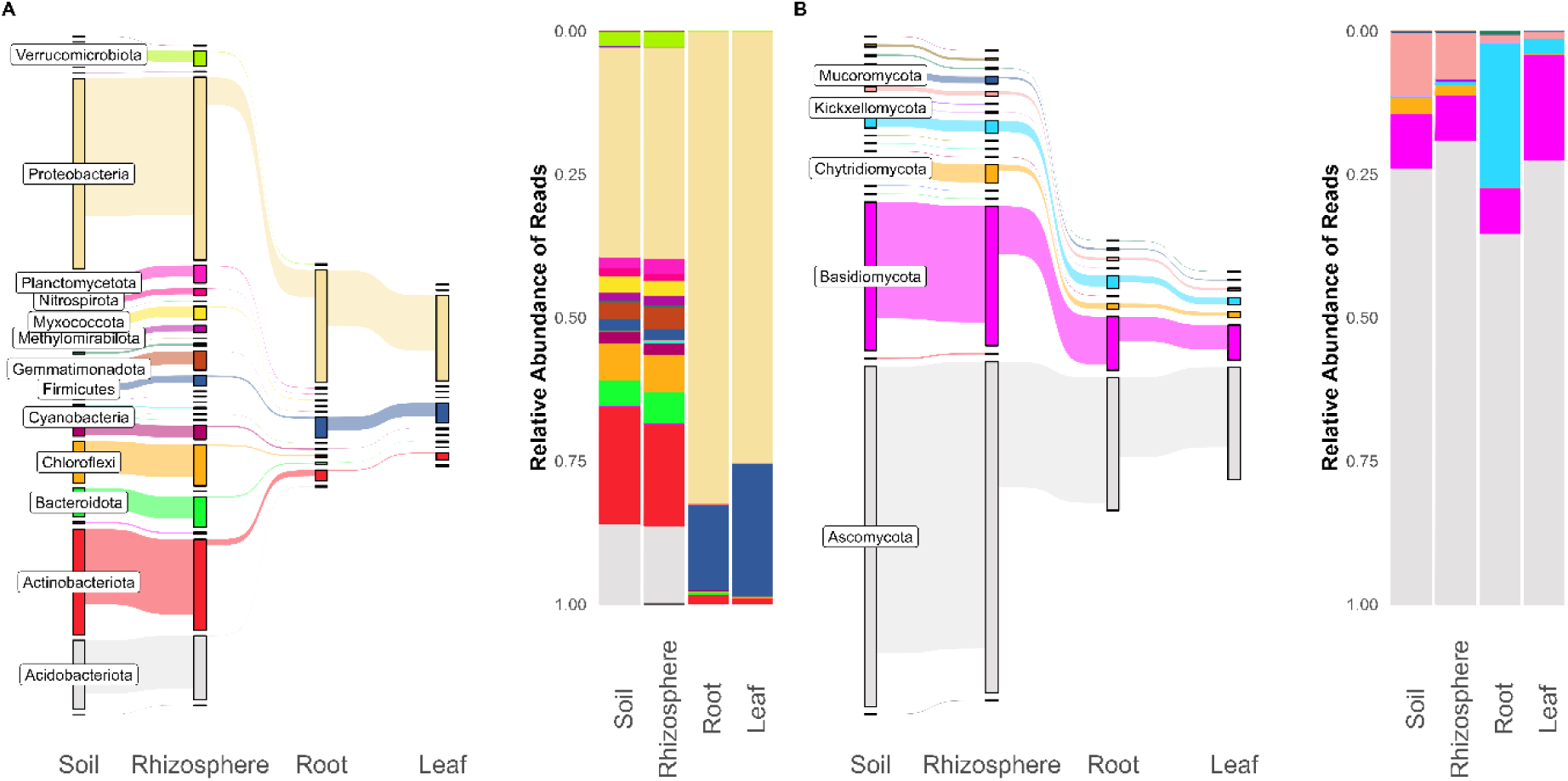
Shared ASVs across plant tissues compared to the relative abundance of reads in plant tissues in (A) bacteria and (B) fungi. Segments are colored by phylum.

ASV data was transformed into proportions (relative abundance) at a read depth of 10,000 reads per sample. Alpha diversity was measured with three common metrics, Observed ASVs, Shannon entropy, and Simpson entropy (Supp Figure 1). Comparisons were made using a marginal PERMANOVA analysis (Supp Table 1). Across all samples we found that bacteria and fungi diversity was chiefly affected by both tissue type and location (Field). When looking just within individual tissue types, bacteria and fungi show opposite effects. Both show a significant association with sample location and not with plant growth stage, but for Bacteria this is only in the soil and rhizosphere, while for fungi it is only in roots and leaves.

We calculated Beta diversity using the weighted UniFrac and Bray Curtis metrics for Bacteria, and Bray Curtis for Fungi (Figure 2). Samples segregated strongly due to tissue type, with less notable segregation based on location (Supp Figure 2). Type II PERMANOVA of weighted UniFrac distances (bacteria) and Bray Curtis distances (fungi) showed that tissue and location had a significant impact on both bacterial and fungal Beta Diversity (p < .01) (Supp Table 2). We compared Bray-Curtis distances and geographic distances between locations using a mantel test, and found that Soil and Rhizosphere Bray-Curtis distances correlated with geographic distance between fields (p < 05) (Supp Table 3).

**Figure 2.**
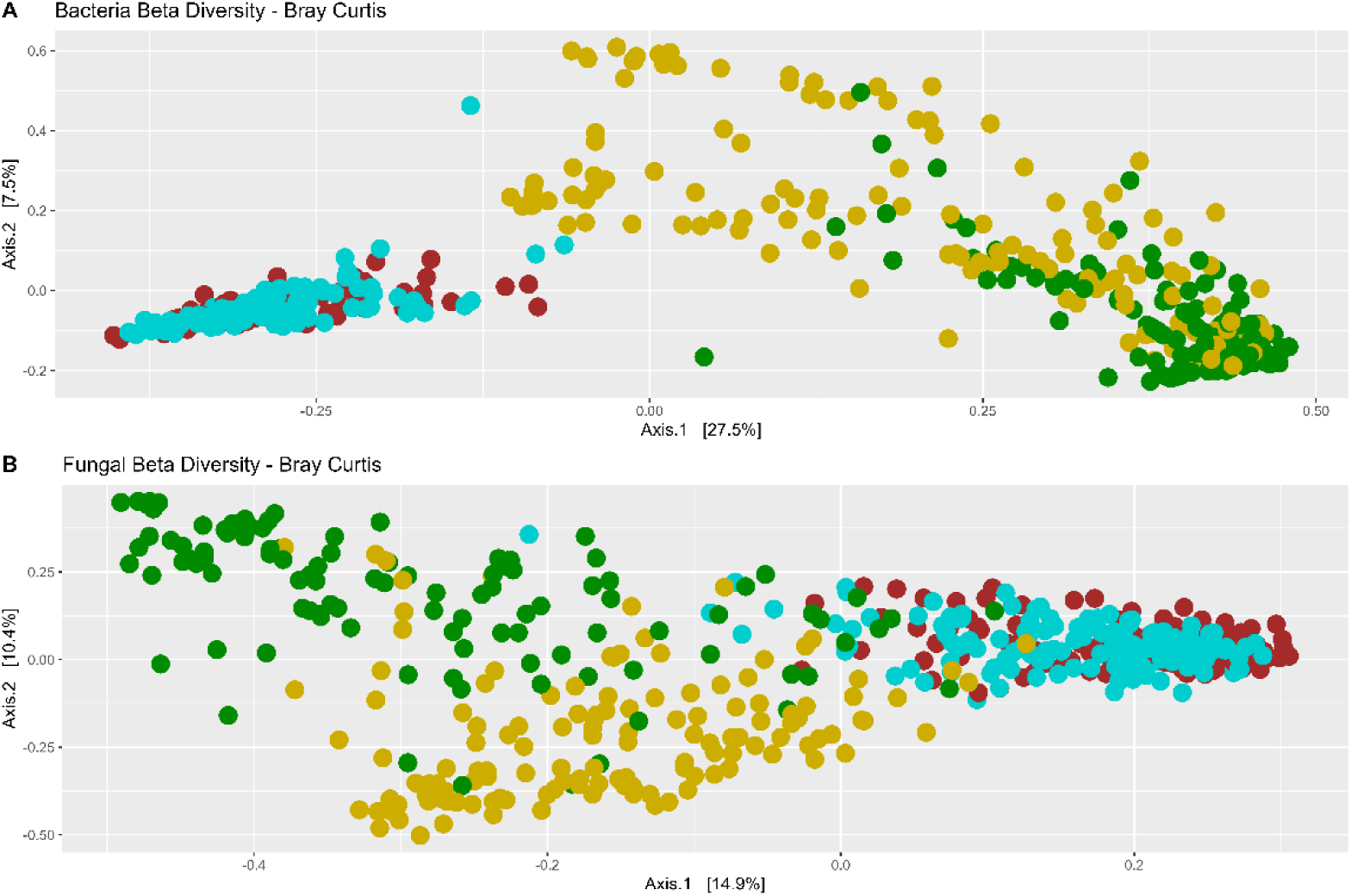
Bray-Curtis (bacteria, top), (fungi - bottom) diversity principle coordinates colorized by tissue type.

We looked to see if differences in environmental and soil measures had an impact on microbial abundance. We used a subset of 2/3 of our samples, as 10 locations were missing soil data. Figure 3 shows the impact environmental covariates have on a number of bacteria and fungi taxa at the phylum level, as this taxa level was the most impacted by environmental factors. Supp Table 4 contains all significant correlations at the phylum level. Overall environmental variables appeared to have impacted taxa in the soil and rhizosphere more than the roots and leaves, and environmental variables affected more fungi taxa than bacteria.

**Figure 3.**
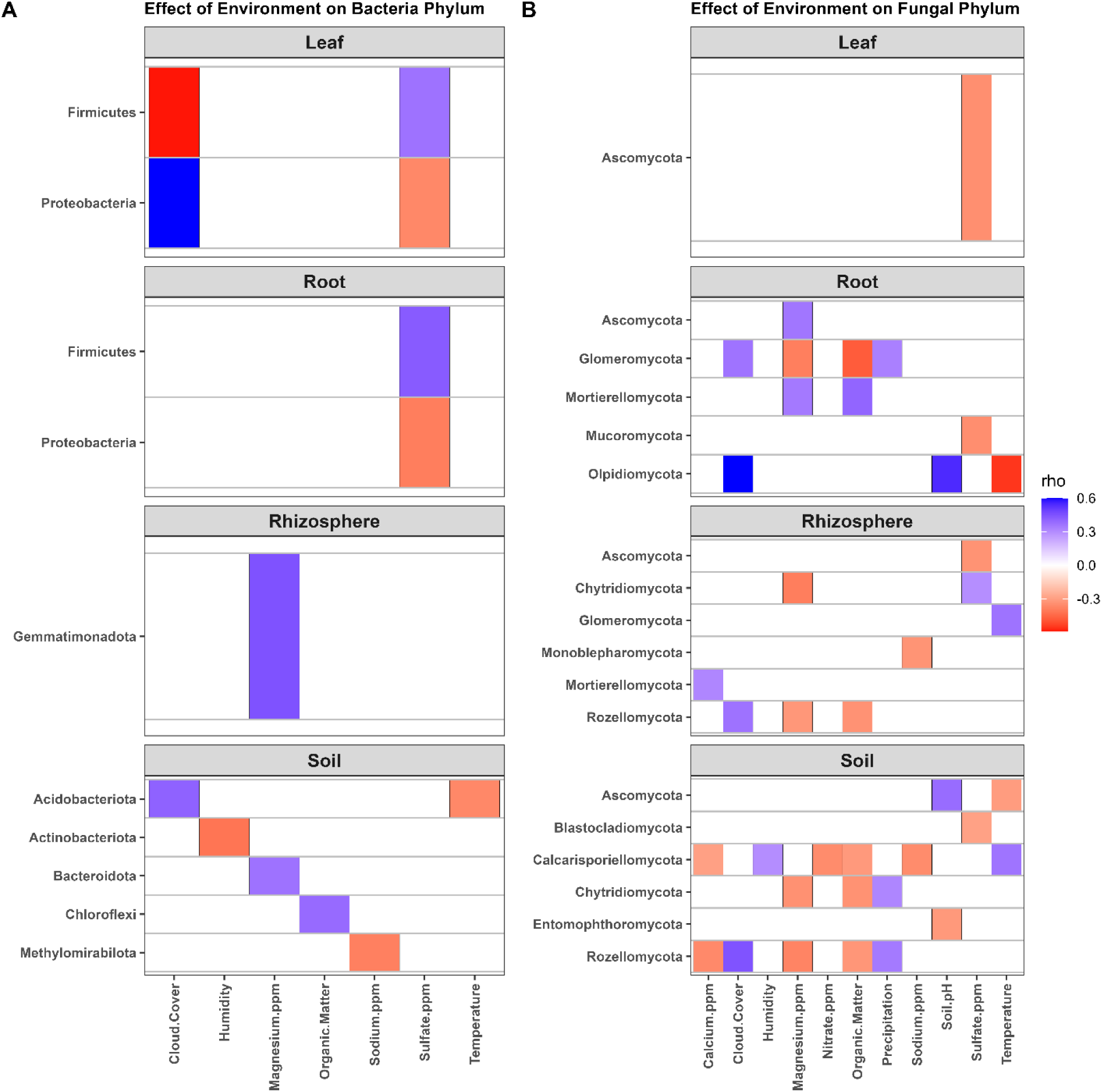
Correlation of bacterial (A) and fungal (B) phyla with environmental parameters. Phylums (Y axis) are significantly correlated with weather and soil covariates (X axis) when there is a colored block present. The color of the block indicates the rho (correlation), showing if the correlation is positive or negative.

### Network Analysis

We then looked at the natural connectivity of networks at the ASV level (Figure 4). Natural Connectivity is a measure of network robustness, and to capture a more granular view we created networks of a single tissue type in a single location using a less strict rho cutoff of 0.3. A Type II ANOVA showed that network natural connectivity was dependent upon tissue type, but not location or the number of taxa in the network (Supp Table 5). We found that leaves and roots had higher natural connectivity than the rhizosphere and soil. While Figure 4 shows natural connectivity at the ASV level, this pattern is consistent across taxa level. Supp Figure 4 shows this analysis at the Phylum level. When we used ANOVAs to see if environmental covariates had an impact on tissue specific network connectivity, the only significant relationships were organic matter and calcium concentration on soil samples (Supp Table 4).

**Figure 4.**
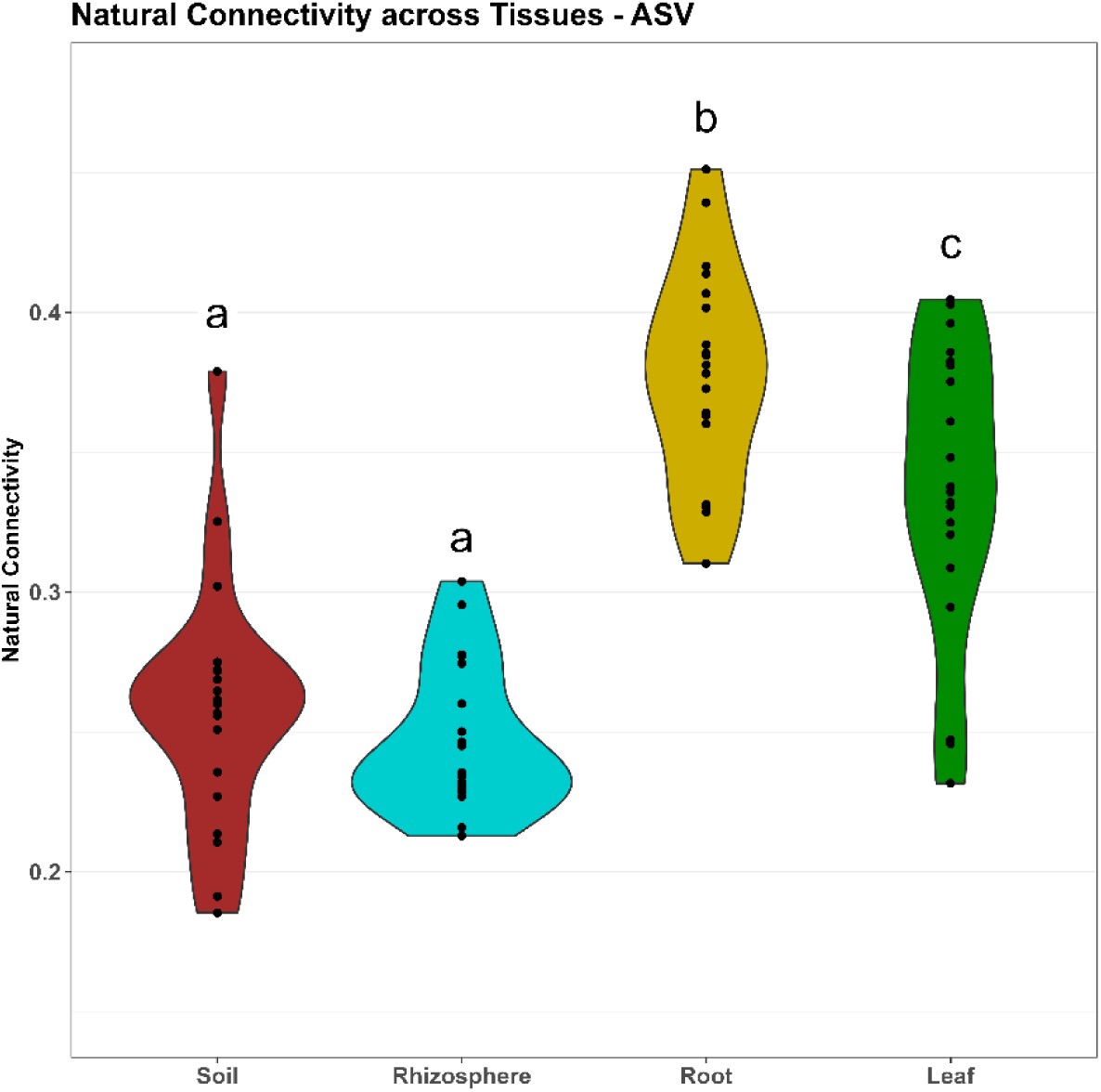
Natural connectivity of samples in a single location, subset by tissue type.

We next constructed networks of our four sample types at the phylum level to investigate potential antagonistic and mutualistic relationships between different bacteria and fungi taxa (Figure 5). Taxonomy was agglomerated at the phylum level, as networks at lower levels are too large and complicated to easily draw conclusions about specific taxa. In order to identify only the strongest relationships, we used a conservative approach with a rho (correlation coefficient) cutoff of 0.8. Across all four sample types, we see bacteria and fungi positively correlated with central hub taxa, which are fungi in the rhizosphere, roots, and leaves. Actinobacteria, Basidomycota, and Mucoromycota were negativily correlated with these central taxa. The archaea phylum Crenarchaeota is positivily correlated with the bacteria Verrucomicrobiota in the roots and leaves. There are more significant correlations in endophyte networks than exterior microbiome networks.

**Figure 5.**
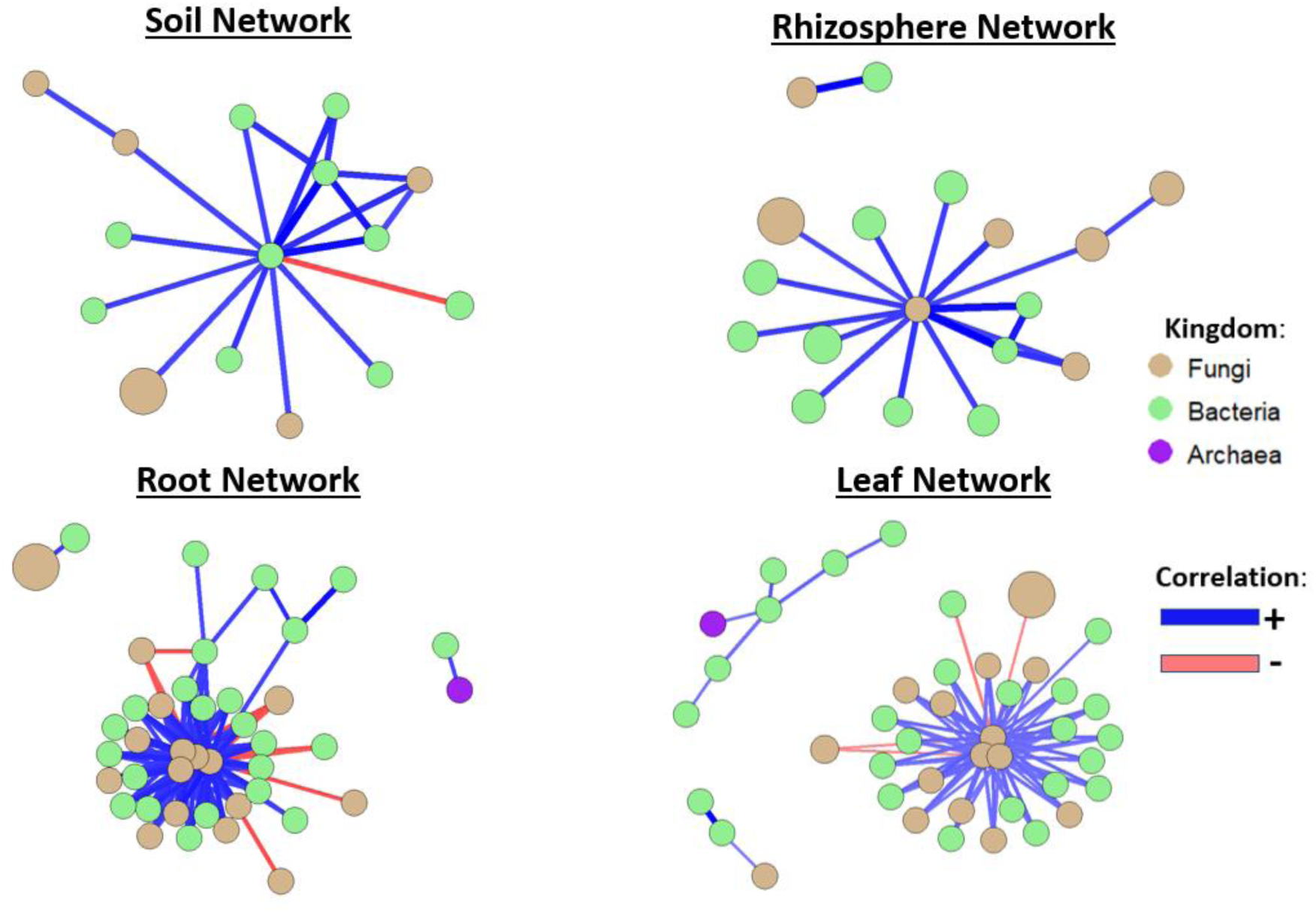
Network of strongly correlated phylums in the soil (A), rhizosphere (B), roots (c) and leaves (d). Edge thickness indicates correlation strength. Hub nodes are labelled with their phyla; a fully labeled version of this figure is in Supp Fig 2.

### Imputed Metagenomics

We used PICRUST2 [101] to predict community core functional gene pathways from 16s and ITS sequences. For the heatmap, these gene pathways were normalized with a variance-stabilizing transformation, and were then clustered by samples (Figure 6) to visualize differences in functional gene profiles. For both bacteria and fungi functional profiles, we saw samples cluster into two groups: exterior organisms and endophytes. These two groups were more similar to each other, and show that there are distinct differences between core microbiome functions based on niche. As shown above, microbial communities of the exterior and interior are more similar to themselves, so the clustering of imputed metagenomics may be a reflection this. These transformed values were turned into a Bray Curtis distance matrix, and a PERMANOVA showed that both tissue and location had significant effects on functional gene composition (p < .001). We then used DESeq2 [104] to identify differentially abundant pathways between individual tissues, as well as a single tissue compared to the other three (Table 1). We found large differences in all comparisons, with one exception: Across all locations soil and rhizosphere core gene functions are almost identical.

**Figure 6.**
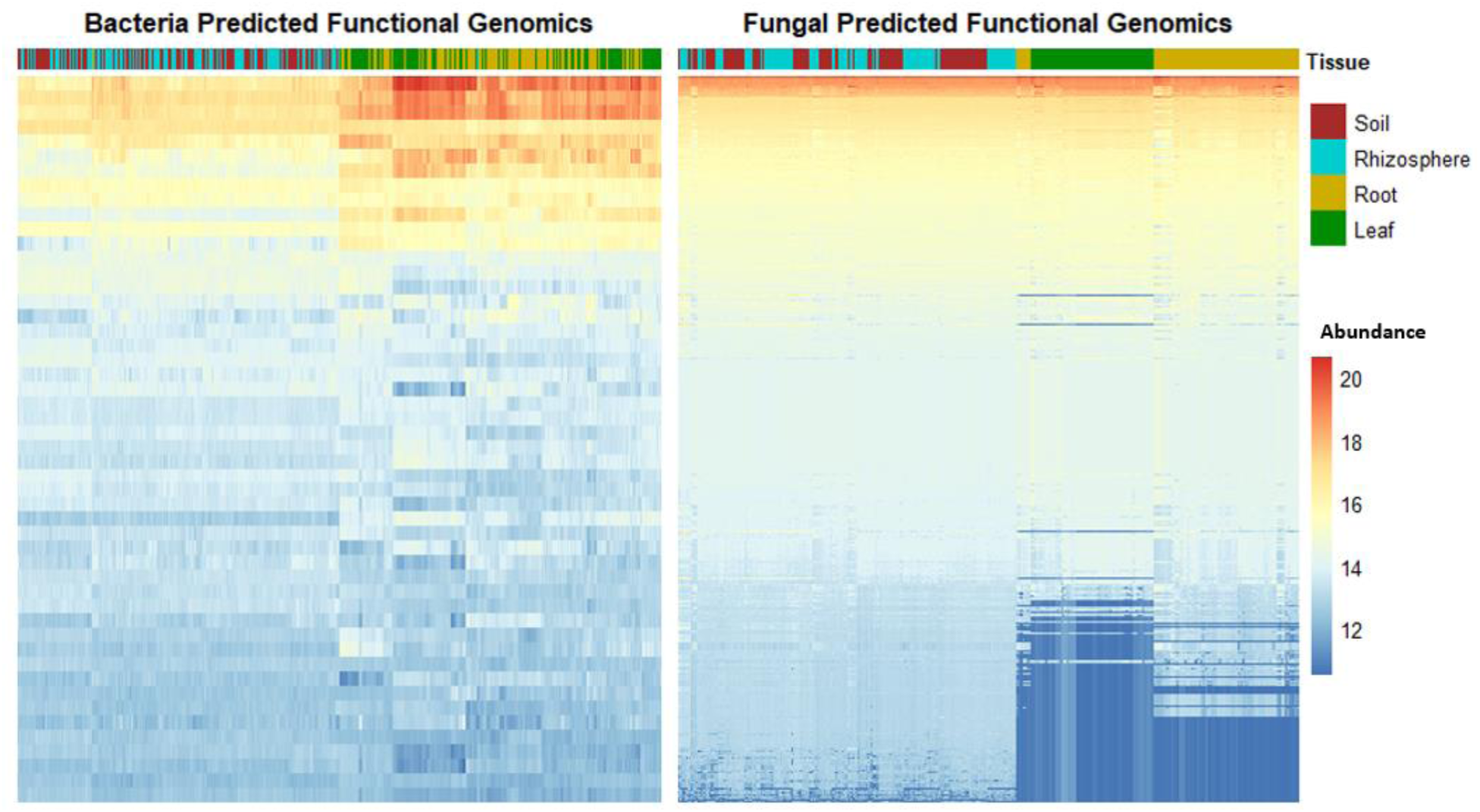
Heat maps of functional profiles for each sample. Samples are clustered based on similarity, and predicted genes are colored by high (red) or low (blue) abundance.

**Table 1.**
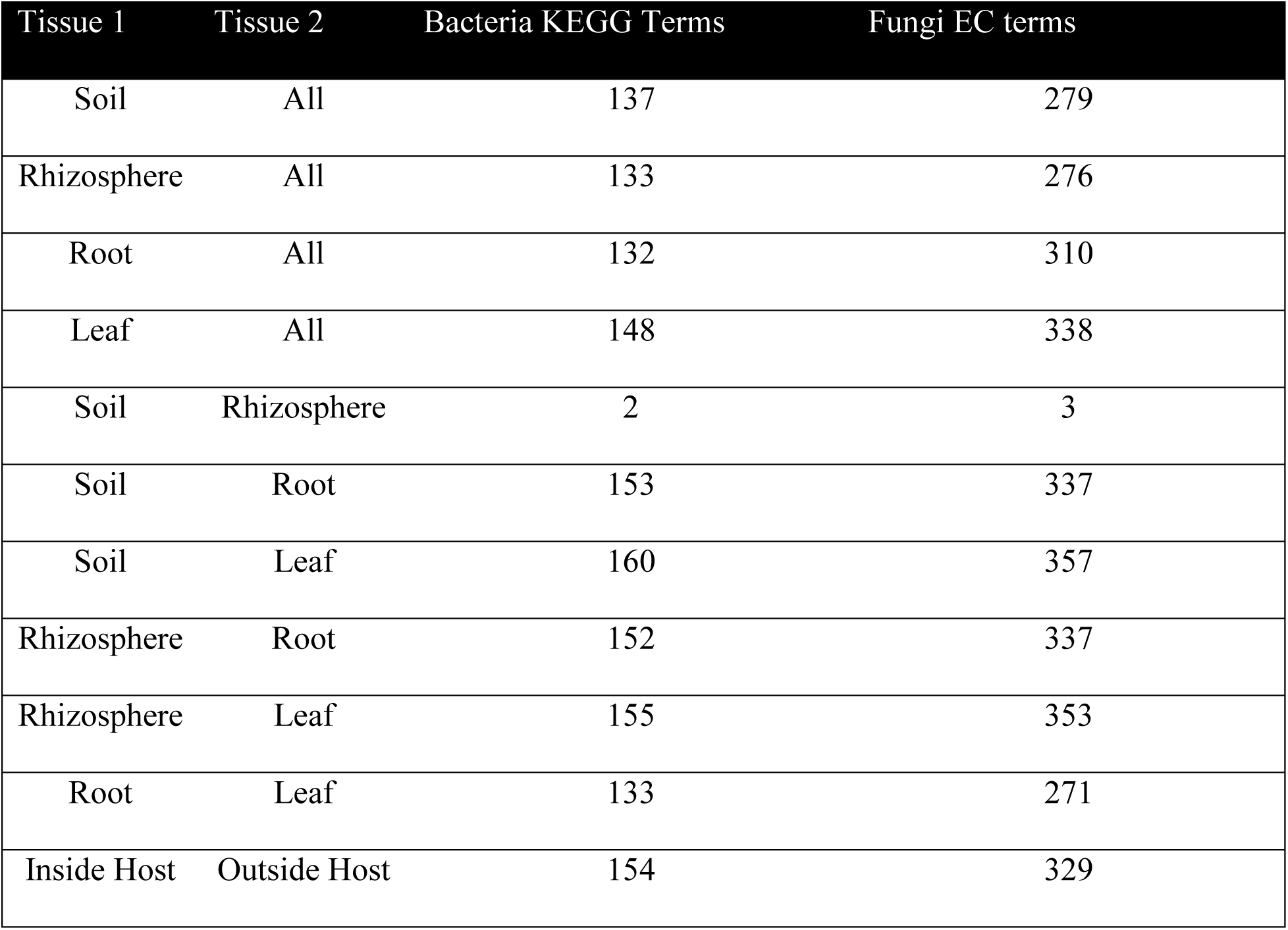
Number of differentially abundant core predicted gene functions by tissue.

| Tissue 1 | Tissue 2 | Bacteria KEGG Terms | Fungi EC terms |
| --- | --- | --- | --- |
| Soil | All | 137 | 279 |
| Rhizosphere | All | 133 | 276 |
| Root | All | 132 | 310 |
| Leaf | All | 148 | 338 |
| Soil | Rhizosphere | 2 | 3 |
| Soil | Root | 153 | 337 |
| Soil | Leaf | 160 | 357 |
| Rhizosphere | Root | 152 | 337 |
| Rhizosphere | Leaf | 155 | 353 |
| Root | Leaf | 133 | 271 |
| Inside Host | Outside Host | 154 | 329 |

We wanted to explore the functional differences of microbes at three distinct transitions: (1) the difference between microbes in bulk soil, and those found in the rhizosphere; (2) the transition of microbes from exterior to endophytes; (3) changes in gene functions in endophytes found in the roots, and those found in the leaves. Supp Data 1 contains all of the differentially abundant comparisons.

Bacteria in the soil had an increase in sporulation gene pathways compared to their counterparts in the rhizosphere. Rhizosphere associated bacteria had increases in hormone biosynthesis. Soil associated fungi had increases in: Nucleoside-triphosphate—adenylate kinase and Magnesium-importing ATPase. Rhizosphere associated fungi had an increase in Farnesol dehydrogenase. Exterior bacteria had distinct differences from bacteria endophytes in the roots and leaves. Bacteria outside the plant had the largest differences in gene pathways related to: biosynthesis of carotenoids, phytohormones, flavonoids, and other potential volatile organic compounds (VOCs); degradation of atrazine; and pathways related to proteasomes and lysosome. Bacteria endophytes had increases in pathways related to: biosynthesis of peptides, bacterial toxins, metabolism and digestion of: carbohydrates amino acids, lipids; mineral absorption, electron transfer carriers, and increases in flagellar assembly. Fungi’s strongest differences were in specific dehydrogenases inside and outside of the plant.

Root bacteria endophytes were enriched for specialized metabolism of VOC pathways including benzoids, phenylpropanoids, and flavonoids. They also have an increased production of streptomycin. Leaf bacteria endophytes had increases energy metabolism such as: Ribosome pathways, electron transfer carriers; amino acid, carbohydrate, and ether lipid metabolism; and the biosynthesis of steroids and carotenoids. Root fungi had large increases in pathways related to membrane transportation: Xenobiotic-transporting ATPase, glyoxylate reductase, molybdate-transporting ATPase, and ethanolamine kinase. Leaf fungi had increases in: farnesol dehydrogenase, sulfatases, and pathways related to carbohydrate and sugar metabolism

## Discussion

Our results show that tissue type and the environment have significant and strong effects on both bacterial and fungal communities in maize. While previous work has shown that host genotype affects microbiome structure and function [33–36], we were unable to take the genetics of our maize into account in this paper. The genetics of this commercial maize are proprietary information, and in several cases, we were unable to positively identify which variety was used. We show that the exterior microbial community is more similar to itself than the endophyte community (Figure 1), and we see a steep drop off of the number of taxa found in the Endosphere of the roots and leaves. We may be able to attribute this to barriers microbes face when colonizing the interior of the plant, as its been shown that endophytes have genetic adaptions compared to exterior brethren [105–108], and several studies have provided evidence of this steep drop off in taxa between the exterior of the plant and the interior [34, 36, 109, 110]. We found that plant tissue and field location both had a statistically significant impact on alpha diversity, with larger separation due to tissue rather than location (Supp Figure 1). When looking within tissue types, we found distinct differences in bacterial and fungal microbiomes. Bacteria communities outside the plant were dependent upon location, as were fungi communities inside the plant. Tissue and location also had significant effect on beta diversity across all samples.

Bray-Curtis distance matrices for both bacteria and fungi in the soil and rhizosphere were correlated with actual distance between sites, indicating that microbial communities inside the plant are not dependent soley on biogeography. Throughout the literature we find that tissue type and the environment have a strong impact on microbiome assembly [3, 33, 34, 36, 67, 111], however, this is the first large scale study with the power to show that tissue compartments can be more similar across the country, than they are to other compartments in the same field.

Fungi appear to be more sensitive to environmental changes, as there were far more interactions between fungi and the environment, specifically in the soil, rhizosphere, and roots. We also see the same group of fungal taxa correlated with different environmental covariates in different tissue compartments. For instance, Rozellomycota, which lack chitinous cell walls, are consistently correlated with cloud cover, magnesium, and organic matter in both the soil and rhizosphere. Yet, it appears that the transition from soil to rhizosphere may alleviate some of the effects that calcium and precipitation have on this taxa. Glomeromycota is only correlated with temperature in the soil samples, yet as a root endophyte it is correlated with cloud cover, magnesium, organic matter, and precipitation. Ascomycota is negatively correlated with sulfate in the soil, rhizosphere and leaves, but positively correlated with magnesium in the root. These correlations tell a story of tissue-specific interactions with the environment, where different niches can perhaps alleviate or enhance a microbe’s sensitivity to weather and soil variables. In this analysis we consistently see cloud cover correlate with taxa abundance. We assume cloud cover is not directly affecting microbes, but is instead a proxy for some other variable(s). Although it correlates with other covariates like solar radiation, solar energy, and UV index, these variables do not correlate with the same taxa, indicating that there is some other variable or combination of variables that cloud cover is a proxy for. We hypothesize that cloud cover may be a proxy for multiple locations or regions of the country. For example, there is some clustering of individual phylum at the location level (New York fields have similar Firmicutes abundance), and cloud cover correlates well with Fimicutes abundance in these fields.

We then investigated natural connectivity of networks in a single tissue, at a single location, at the ASV level (Figure 4). As a single measure describing the amount of correlation between microbes, we view natural connectivity as a measure of potential interactions. A high natural connectivity may indicate that there could me more real biological interactions. At this granular level, we found that the endosphere has a significantly higher natural connectivity than the soil and rhizosphere. Although the soil and rhizosphere microbiome is larger and more diverse than the endosphere, it appears that endosphere ASVs are more connected with each other. This trend also holds when taxa are agglomerated to the phylum level (Supp Figure 4). This indicates that within a single location, more microorganisms may be interacting with each other inside the plant, perhaps weakly.

We found consistent patterns of inter-kingdom antagonism and intra-kingdom mutualism between bacteria and fungi. Figure 5 demonstrates large patterns of correlations across all 30 locations at the phylum level. Although networks in the soil and rhizosphere were much larger and more complex than in the endosphere, exterior networks were not larger than endopshere networks. Interestingly, in the plant-interior samples the Archaea phylum Crenarchaeota is strongly associated with the bacteria Verrucomicrobiota. Although not often discussed, Archaea have been found to partake in the nitrogen cycle in agricultural soils [112–114], and their interactions with bacteria and fungi provide an interesting area for further research. Across all tissues we see positive correlations between bacteria and fungi with hub taxa, with very few negative correlations. These correlations may inticate interactions in these communities. It’s been shown that bacteria can act as mutualists with mycorrhizal fungi and can increase growth promotion in the host [115, 116], and in a gnotobiotic experiment, it was shown that a nitrogen fixing bacteria and a mycorrhizal fungi acted as mutualists to increase growth promotion and drastically alter gene expression in the host plant and fungus [117]. The literature frequently deomonstrates antagonism between bacteria and fungi [118]. It has been shown that fungi often secrete anti-biotics [119, 120], and can disrupt bacteria communication in forest soils [121]. Soil bacteria can suppress fungi through volatile secretions [122], and one study showed that increases in fungal density was associated with enrichment of bacteria that possessed antifungal mechanisms, such as siderophores, cyanide, and lytic enzymes [56]. These correlations can only hint at potential interactions, interactions that may be indirect (where environmental factors regulate their abundance such as Figure 3), or direct interaction between these organisms.

Further studies of maize tissue microenvironments could elucidate what is causing these differences by examining resource availability, host defense stressors, and quantifying levels of molecular antagonism amongst the community in each compartment. Our inferred metagenomics analysis found community differences that support differences in antagonism throughout maize tissue compartments.

We wanted to explore the functional differences of microbes at three distinct transitions: (1) the difference between microbes in bulk soil, and those found in the rhizosphere; (2) the transition of microbes from outside the plant to within; (3) changes in gene functions in endophytes found in the roots, and those found in the leaves. Supp Data 1 contains all of the differentially abundant comparisons. Our functional gene profiles showed clear and distinct clustering of samples from within or outside plant tissue (Figure 6.) which is consistent with the rest of our findings. When looking for differentially abundant core pathways, we found massive differences in gene profiles for both bacterial and fungal communities (Table 1); the only exception was comparing soil to rhizosphere.

Bacteria in the soil had an increase in sporulation gene pathways compared to their counterparts in the rhizosphere. Sporulation results in metabolically inactive and resistant spores [123], and this process leaves highly resistant dormant spores that can activate when conditions improve. This implies that Bacteria in the rhizosphere become less dormant, perhaps association with maize roots or their exudes can protect bacteria from environmental stressors, just like bacteria can protect maize from abiotic stress [6, 12, 124]. It may also indicate that the soil environment doesn’t favor spore-forming groups in general. Rhizosphere associated bacteria also had increases in hormone biosynthesis. These phytohormones can be beneficial to the plant and are important in microbe-host communication [13–15], and its believed that the host can select for bacteria in the rhizosphere that can promote growth [111]. Rhizosphere associated fungi had an increase in Farnesol dehydrogenase. Farnesol can be used in fungi to fungi communication, as well as an antagonist against bacteria quorum sensing [125]. It’s been shown that bacteria form biofilms on plant roots and this interaction is important for plant growth promotion [126], which could lead to fungi antagonism disrupting this quorum sensing as the fungus also associates with the maize root.

Exterior bacteria were enriched for pathways the can be part of soluble molecule metabolism as well as known VOC metabolism (phenol, terpenes, hydrocarbons) compared to endophytes. The rhizosphere is a hotspot for signaling between the plant host and microbes as reviewed in [15, 127–131]. Root exudes account for almost 10% of photosynthetically fixed carbon and 15% of plant nitrogen [128]. These compounds released from the plant can shape the microbiome, recruit beneficial organism, and can inhibit bacteria. For example it was shown that maize’s production of benzoxazinoids, can shape bacteria and fungi abundance in the rhizosphere [132–134]. Phenolics, flavonoids, and indole-3-acetic acid were crucial in recruiting *Aspergillus nominus* wlg2 to the rhizosphere [135]. We found enriched pathways for compounds from these groups and more, in exterior bacteria from around the country. Endophytes were more mobile, and could metabolize more energy rich compounds. The interior of the plant is energy-rich compared to field soil. It also had a type IV pili, which is important for bacteria motility [105]. A comparison of *Herbaspirillum* species found grass endophytic species had increases in carbohydrate and nitrogen metabolism, and plant cell wall degradation compared to their free-living counterparts [107]. A study of the rice microbiome also found increases motility via flagella, and plant-polymer degrading enzymes [108].

When we compared root endophytes to leaf endophytes, we found potential similarities between bacteria and fungi. Root bacteria were enriched for VOC metabolism (phenols, terpenes, hydrocarbons), and fungi were enriched for enzymes related to membrane transport of inorganic compouns. These mechanisms may be ways to adapt to, and take advantage of, the large diversity of molecular signaling that occurs between microbes and the host [15, 127, 136]. Both bacteria and fungi in the leaves had increases in pathways related to carbohydrate metabolism and energy pathways. In the leaves these organisms have direct access to photosynthates, which bacteria and fungi use as an energy source [137]. It has been proposed that endophytes play a role in plant photosynthesis through a number of mechanisms, including carbon-dioxide assimilation [137, 138]. In rice, it was estimated that microbial respiration accounted for 57% of carbon-dioxide in the plant [137].

Molecular interactions between bacteria, fungi, and their host, are highly complex and is a blossoming area of research [118, 127, 129, 130, 139, 140]. If we assume our metagenome functions represent an accurate view of the microbes in each compartment, then it implies that there are large shifts in community function during key transitions: (1) Bacteria have more genes related to phytohormone production and fungi have more genes related to farnesol dehydrogenase when moving from soil to the root surface. (2) A larger fraction of bacteria outside the plant are capable of degrading and producing more VOC’s than endophytes, and endophytes have the capacity to be more mobile, and have the capacity to take advantage of more available resources. Fungi dehydrogenase diversity show endophytes have genes capable of metabolizing different plant-exudes compared to their soil and rhizosphere associated brethren. (3) More root bacteria can metabolize VOC’s than leaf bacteria, and leaf bacteria overall have more genes related to energy pathways. Root fungi have more pathways related to membrane transport, while leaf fungi have the more genes related to metabolizing carbohydrates from the plant and increases in farnesol related genes again. In every comparison, we see differences in metabolites that may be involved with chemical antagonism, whether it be defense mechanisms from the host, antibiotics, or potential anti-fungals. Previous work has shown that predicted metabolic functional pathways correlate with actual metabolomics data in support of PICRUST2 predictions [141]. However, it must be noted that ITS prediction accuracy is much lower than 16s prediction accuracy [101].

In this study we sought to (1) characterize bacterial and fungal communities, (2) identify inter-kingdom interactions (3) identify the impact environment has on microbiome networks, and (4) identify the core functions of the commercial maize microbiome. We found that (1) communities of bacteria and fungi were more strongly influenced by tissue than environment, and that across the US these communities in the soil, rhizosphere, roots, and leaves are dominated by Proteobacteria and Ascomycota; (2) we found patterns of potential interactions between fungi and bacteria in all four tissues, and found that endosphere networks are more tightly connected; (3) we found that the environment effected individual taxa, community structure, and function, but did not affect the natural connectivity of networks; (4) we found tissue type and location significantly affect predicted gene function profiles, and that the functional pathways of communities vary to deal with very specific environments.

## Conclusion

The maize microbiome is well studied, and understanding the interactions between host and microbial community is crucial to make further strides in crop protection and enhancement. Prior research has shown that the host and environment play a large role in microbial community assembly. In this paper we examined the structure, potential function, and interactions of maize bacteria and fungi communities across 30 locations in the United States. We found tissue niche had a larger effect on the maize microbiome than location. Tissue compartments around the country are structurally and functionally more similar to each other than to other tissues in the same field. Throughout our analysis, we find distinct differences between exterior microbes versus endophytes, including predicted gene functions that may be critical for endophyte colonization. This work has proposed that there are distinct differences in the general functions of tissue-specific microbiomes, but further projects are necessary to validate them. In the future, controlled environment studies, synthetic communities, and meta-transcriptomics could help elucidate exactly how these interactions shape plant host growth and health.

## Supporting information

Supplemental Figs and Files zip

## Supplementary Materials

SF1 Alpha Diversity Figure

ST1 Alpha Diversity Stats

SF2 Beta by Location

ST2 Beta Diversity PERMANOVA

ST3 Mantel Test

ST4 Env Corr DFs

SF3 Labeled Networks

SF4 Phylum Natural Connectivity

ST5 Natural Connectivity Soil Permanova

SD1 Folder containing PICRUST analysis.

## Author Contributions

Conceptualization, J.G.W. and C.R.S.; methodology, C.R.S. formal analysis, C.R.S and H.D.; writing—original draft preparation, C.R.S.; writing—review and editing, J.G.W.; visualization, C.R.S.; supervision, J.G.W.; project administration, J.G.W.; funding acquisition, J.G.W. All authors have read and agreed to the published version of the manuscript.

## Data Availability

The Genomes to Fields and Indigo Ag Microbiome Data 2017 is publicly available and was downloaded from Cyverse data commons (DOI: 10.25739/htck-sw56). Weather, soil, and field meta data was also downloaded from Cyverse data commons (DOI:10.25739/frmv-wj25). All bioinformatics scripts and pipelines are available at https://github.com/wallace/lab/paper-schultz-indigo-microbiome-2023.

## Acknowledgments

We would like to acknowledge all of the participants of the Genomes to Fields initiative and Indigo Ag for making this data publically available.

