## Supplementary figures and images for "The Landscape of Maize-Associated Bacteria and Fungi Across the United States"

### SF1_Bacteria_AlphaDiv.png

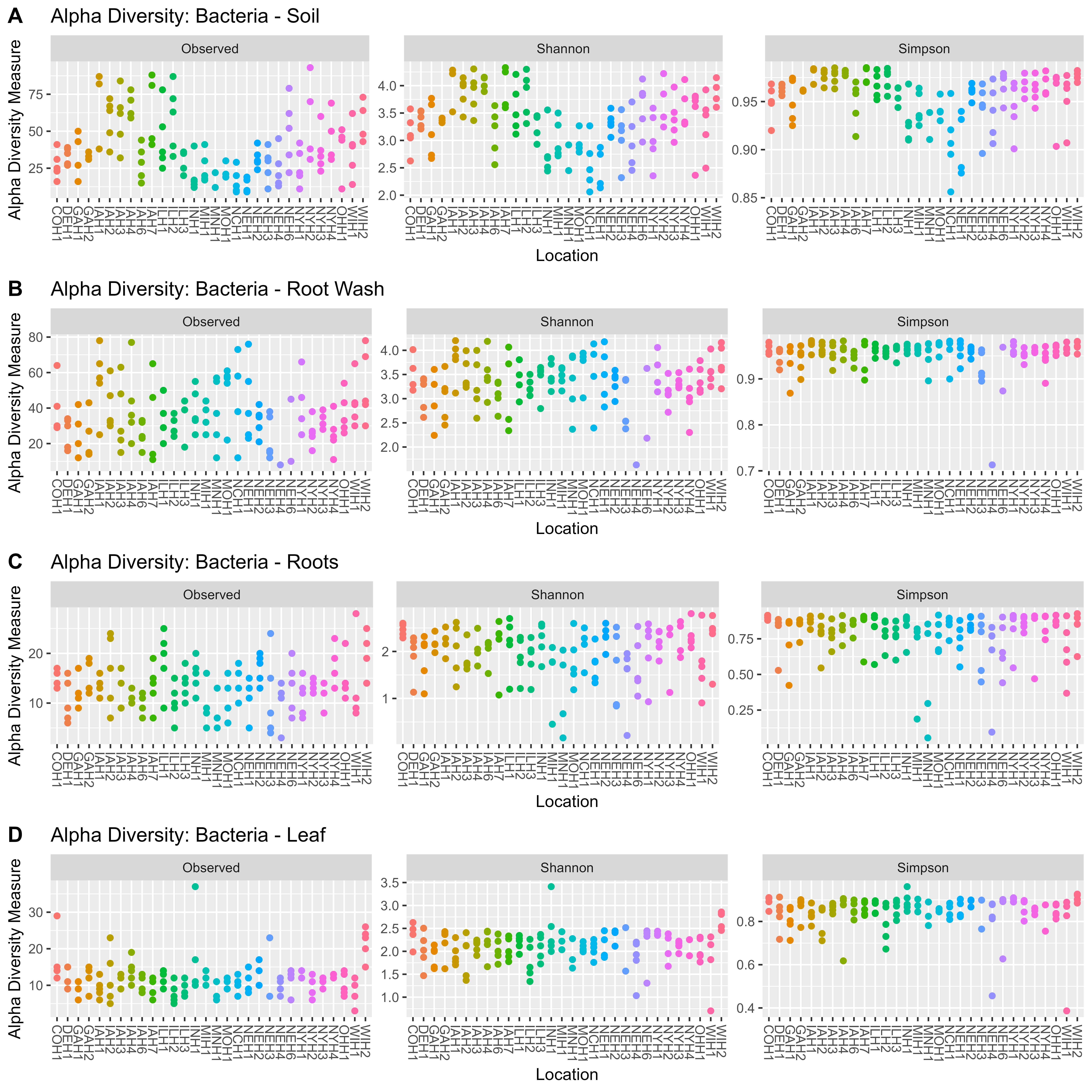

### SF1_Fungi_AlphaDiv.png

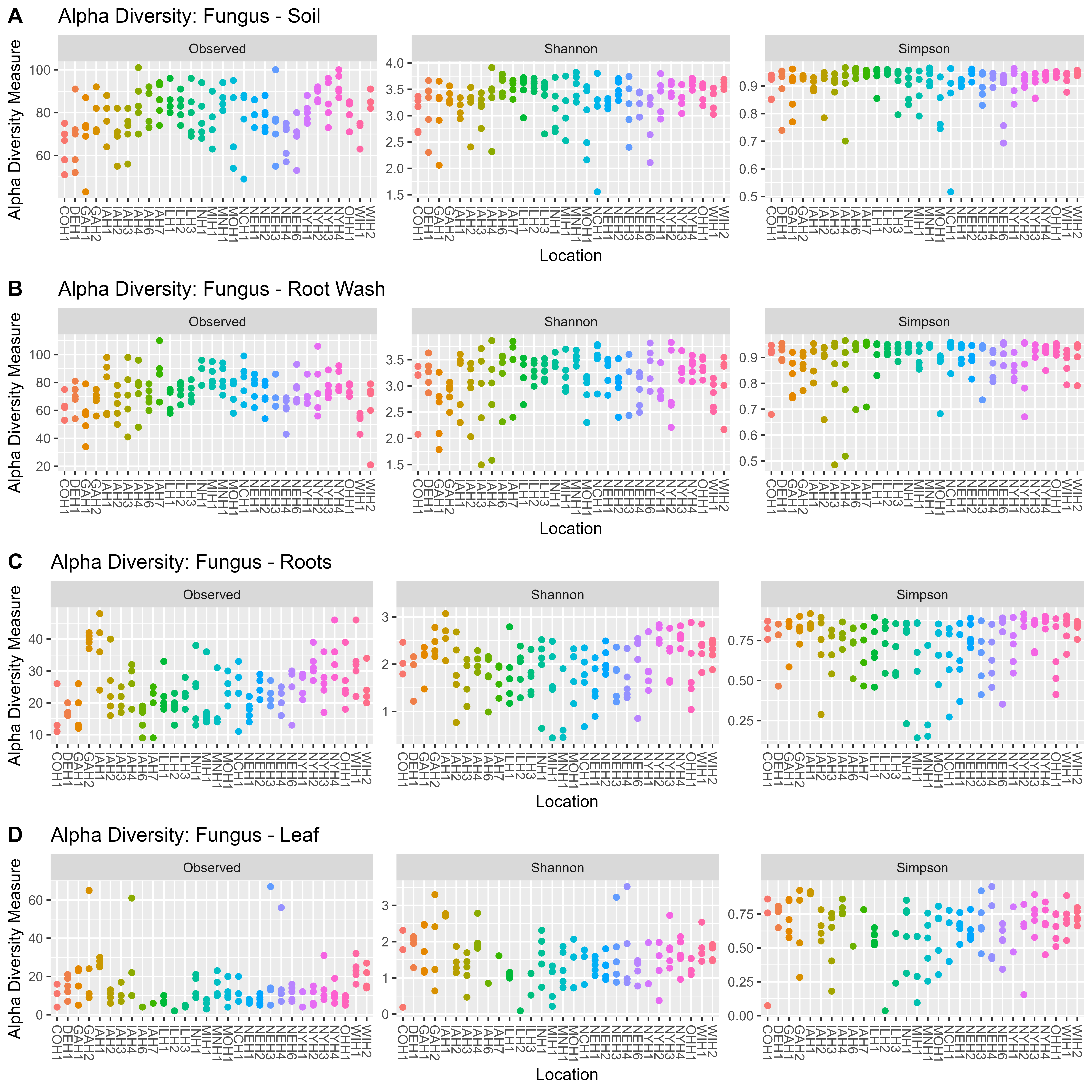

### SF2_BetaDiv_Location.png

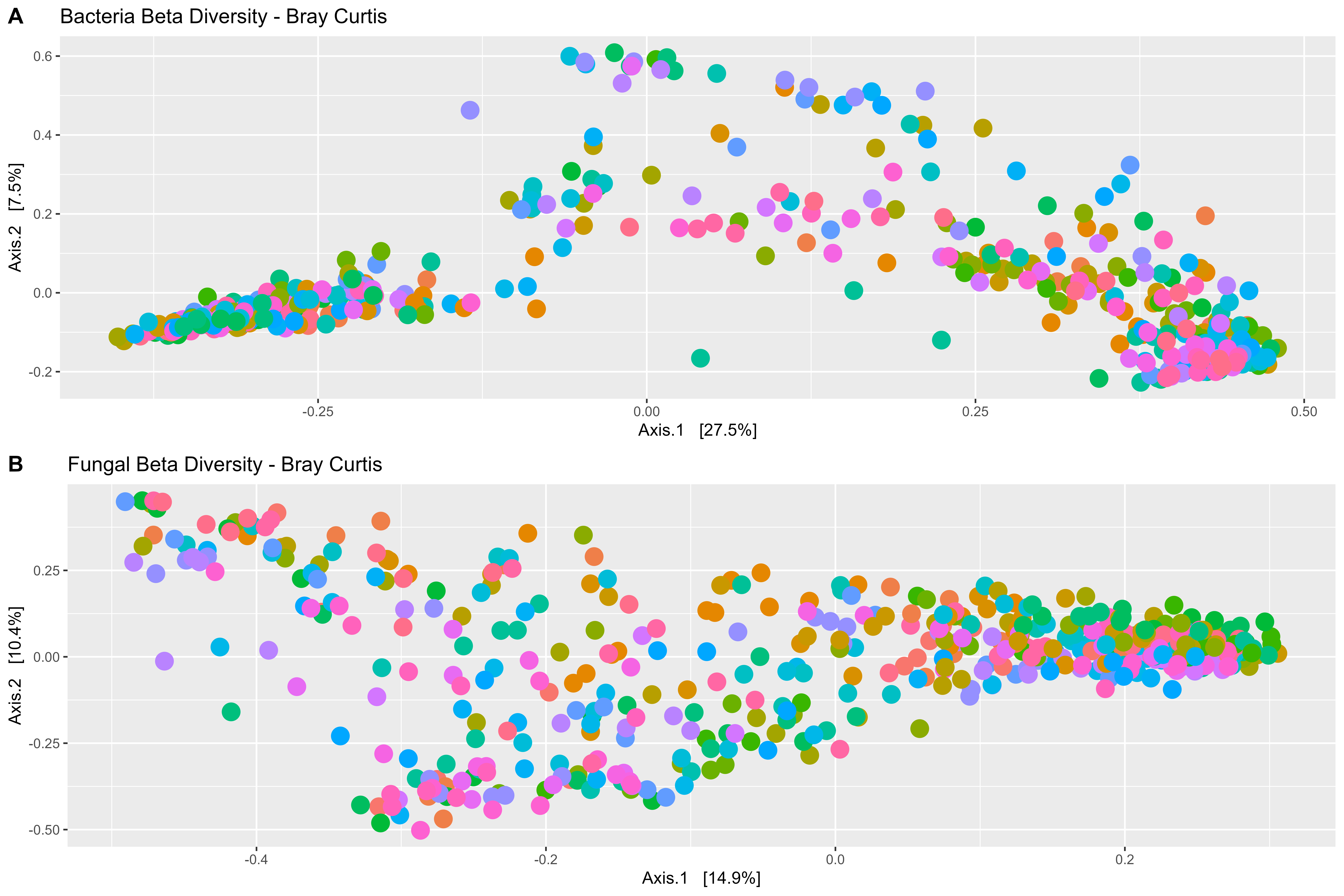

### SF3_Leaf.png

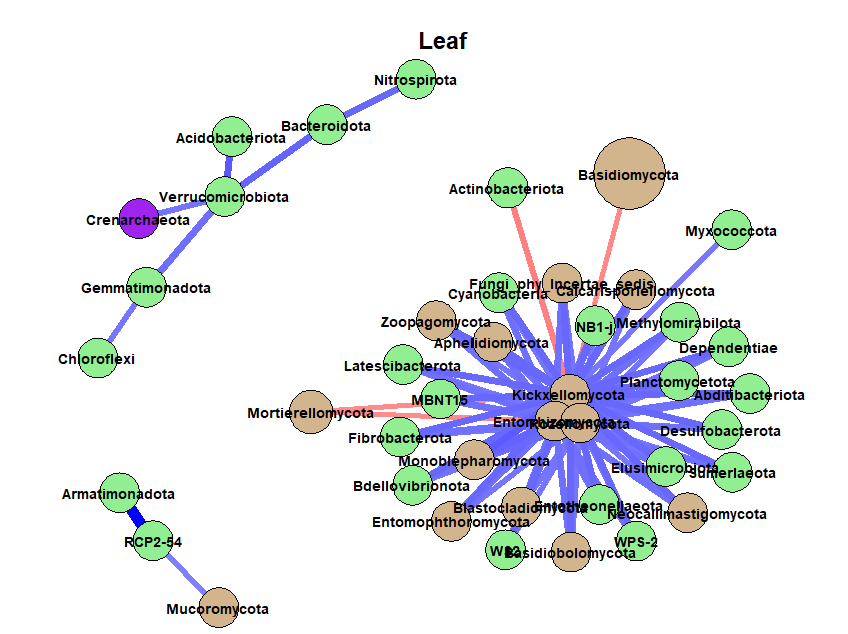

### SF3_Rhiz.png

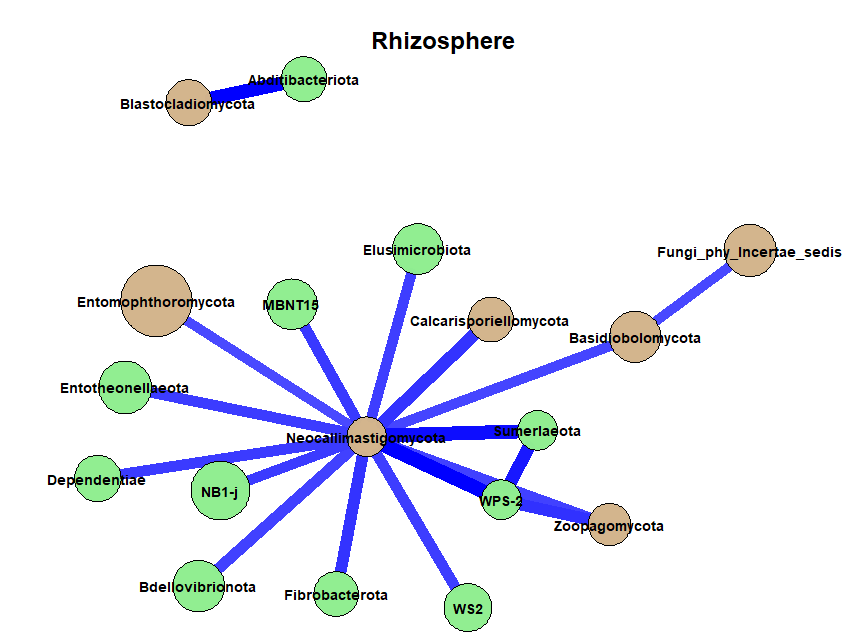

### SF3_Root.png

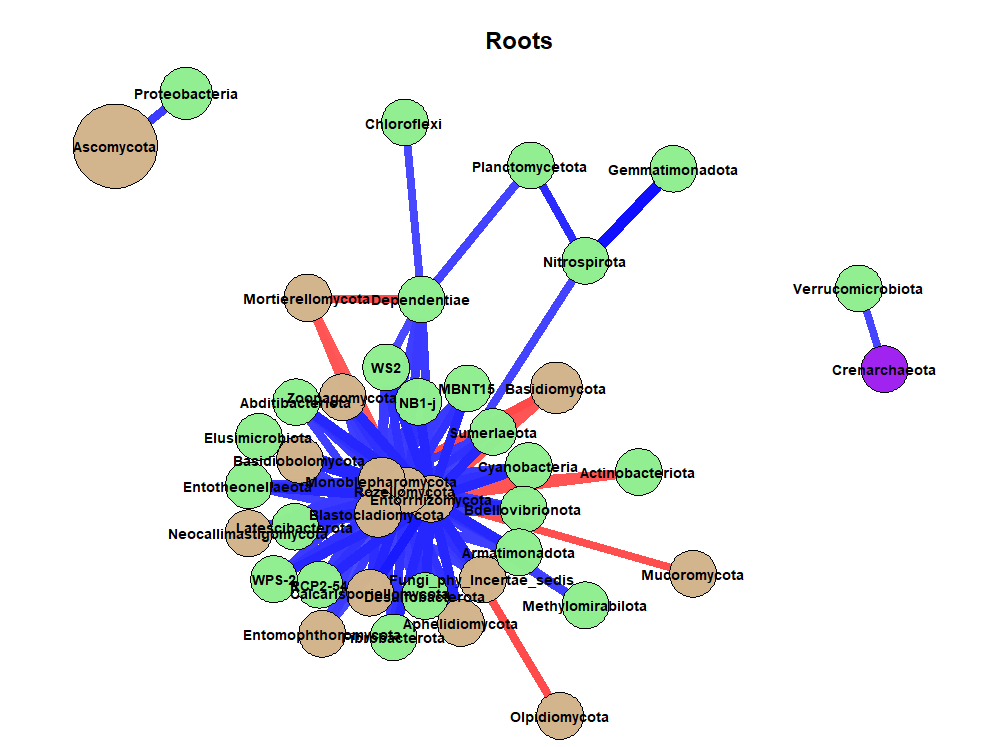

### SF3_Soil.png

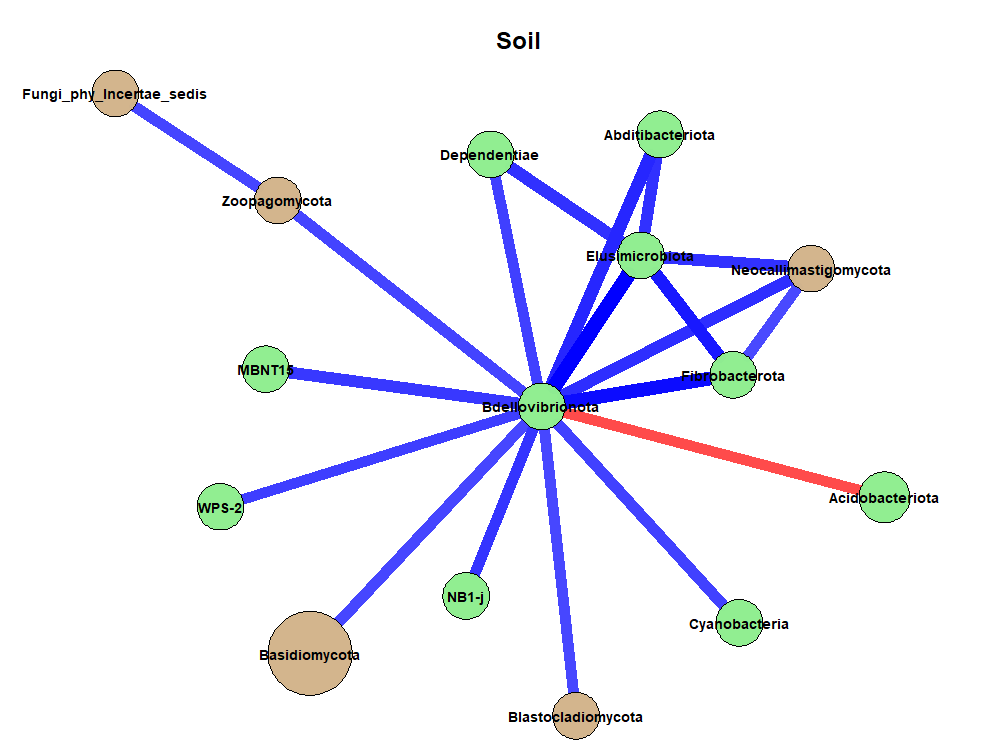

### SF4_NatConnect_Phylum.png

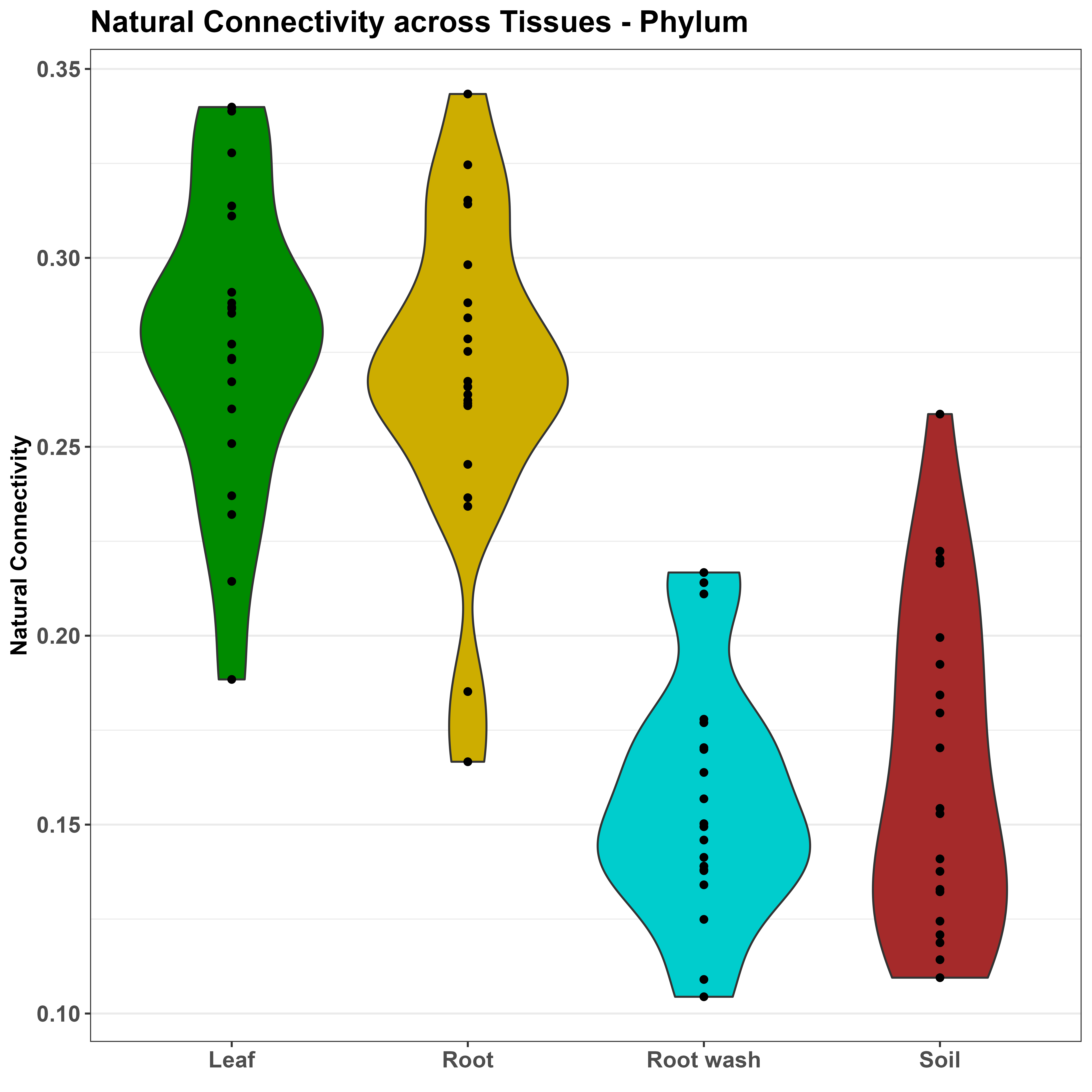
